# Expression of SSEA4 and TRA1-60 as Marker of Induced Pluripotent Stem Cells by Small Molecule Compound VC6TFZ on Peripheral Blood Mononuclear Cell

**DOI:** 10.1101/2020.12.13.422599

**Authors:** A Andrianto, Adityo Basworo, Ivana Purnama Dewi, Budi Susetio Pikir

## Abstract

**Introduction:** It is possible to induce pluripotent stem cells from somatic cells, offering an infinite cell resource with the potential for disease research and use in regenerative medicine. Due to ease of accessibility, minimum invasive treatment, and can be kept frozen, peripheral blood mononuclear cells (PBMC) were an attractive source cell. VC6TFZ, a small molecule compound, has been successfully reprogrammed from mouse fibroblast induced pluripotent stem cells (iPSCs). However, it has not been confirmed in humans.

**Objective:** The aim of this research is to determine whether the small molecule compound VC6TFZ can induced pluripotency of PBMC to generate iPSCs detected with expression of SSEA4 and TRA1-60.

**Methods:** Using the centrifugation gradient density process, mononuclear cells were separated from peripheral venous blood. Mononuclear cells were cultured for 6 days in the expansion medium. The cells were divided into four groups; group 1 (P1), which was not exposed to small molecules (control group) and groups 2-4 (P2-P4), the experimental groups, subjected to various dosages of the small molecule compound VC6TFZ (VPA, CHIR, Tranylcypromine, FSK, Dznep, and TTNPB). The induction of pluripotency using small molecule compound VC6TFZ was completed within 14 days, then for 7 days the medium shifted to 2i medium. iPSCs identification in based on colony morphology and pluripotent gene expression, SSEA4 and TRA1-60 marker, using immunocytochemistry.

**Results:** Colonies appeared on reprogramming process in day 7th. These colonies had round, large, and cobble stone morphology like ESC. Gene expression of SSEA4 and TRA 1-60 increased statisticaly significant than control group (SSEA4 were P2 p=0.007; P3 p=0.001; P4 p=0.009 and TRA 1-60 were P2 p=0.002; P3 p=0.001; P4 p=0.001).

**Conclusion:** Small molecule compound VC6TFZ could induced pluripotency of human PBMC to generate iPSCs. Pluripotency marker gene expression, SSEA 4 and TRA 1-60, in the experimental group was statistically significantly higher than in the control group.

## 1. Introduction

Pluripotent stem cells are expected to become future regenerative therapies that have the ability to differentiate into all cell types derived into three germinal layers. Embryonic stem cells (ESCs) that are produced at the blastocyst stage from the internal cell mass of the embryo are the common source of pluripotent stem cells. The use of ESCs has some of disadvantages due to some ethical issue related to embryo destruction, risk of immune rejection, and limited source due to embryo origin. Induced pluripotent stem cells (iPSCs) have been an alternative source of pluripotent stem cells which tackled the ethical issue because it originates from somatic cells. Potential use of iPSCs such as cells for cell transplantation therapy, diseases modeling, and drug screening [1].

Induced pluripotent stem cells were produced by reprogramming process using different reprogramming factors from somatic cells. The first source cells were skin fibroblast cells, now the most commonly used. [2,3]. Skin biopsy procedures, however have been unconvertable, leave scar tissue, and take a long time for the expansion of fibroblast cells, restricting the use of fibroblasts as iPSC source. Peripheral blood cells have become an attractive source of iPSCs because they are easy to extract, minimally invasive and can be preserved in a frozen state [4].

By transduction of exogenous transcription factors SOX2, OCT4, Klf4, and c-Myc (OSKM) into the nucleus of somatic cells transferred by retroviruses, iPSCs were produced [2,3]. A reprogramming process utilizing exogenous transcription factors with integrative systems that are associated with low efficacy, risk of mutagenesis and teratogenesis, thus restricting their use for clinical applications. To enhance reprogramming efficiency using exogenous transcription factors, small molecules that worked on signaling pathways, epigenetic modifications, and metabolic processes were commonly used. In spite of the lack of exogenous transcription factors [5], the combination of several small molecules may induce pluripotency [5].

VC6TFZ was a combination of small molecules consisting of valproic acid (VP A), CHIR990210 (CHIR), 616452 (Repsox), tranylcypromine, forskolin, 3-deazaneplanocin (DZnep), and 4-[(E)-2-(5,6,7,8-Tetrahydro-5,5,8,8-tetramethyl-2-naphthalenyl)-1- propenylbenzoic acid, or also called TTNPB. In mouse embryonic fibroblasts (MEFs), the combination of these tiny molecules could induce pluripotency with higher efficiency than the Yamanaka protocol that used exogenous transcription factors (0.02 percent vs. 0.001-0.001). [6].

Induced pluripotent stem cells have several heterogeneous characteristics and morphology, so that there are quite a number of markers that can be used to identify the expression of iPSCs. Alkaline phosphatase staining is a marker for surface cell antigen on iPSCs. SSEA4 and TRA 1-60 are antibodies that express the presence of cell surface antigens, particularly against iPSCs. Surface cell antibodies are the most widely used method specifically to identify stem cells in heterogeneous cultures, specifically SSEA4 and TRA 1-60 antibodies present as a benchmark for identifying iPSCs by immunostaining methods. The aim of this research was to identified expression of SSEA4 and TRA1-60 as marker of iPSCs by small-molecule compound VC6TFZ in human peripheral blood mononuclear cells (PBMC).

## 2. Methods

This study is laboratory experimental study (in vitro study) by administrating small molecule compound VC6TFZ to induced pluripotency of PBMC and generate iPSCs. The aim of this research is to determine SSEA4 and TRA1-60 expression as a marker of iPSCs. This study is a true experimental randomized study, with “post-test only control group design” approach (Figure 1). The blood sample for this study derived from a 29-year-old male volunteer. The study protocol, as shown in Figure 2, consisted of several steps including PBMC isolation, PBMC culture, pluripotency induction using small molecules, and iPSCs identification. This research was conducted at the Laboratory of the Center for Research and Development of Stem Cell, Institute of Tropical Disease (ITD), Airlangga University, Surabaya in November 2019 - January 2020.

**Figure 1.**
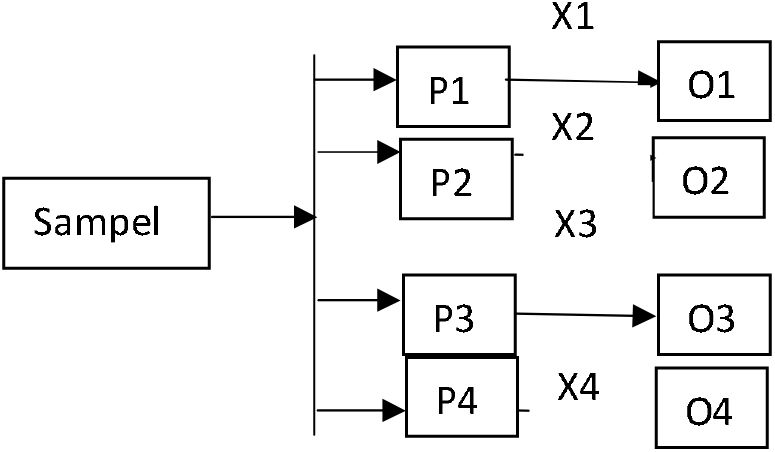
Study Design “Posttest Only Control Group Design”

**Figure 2.**
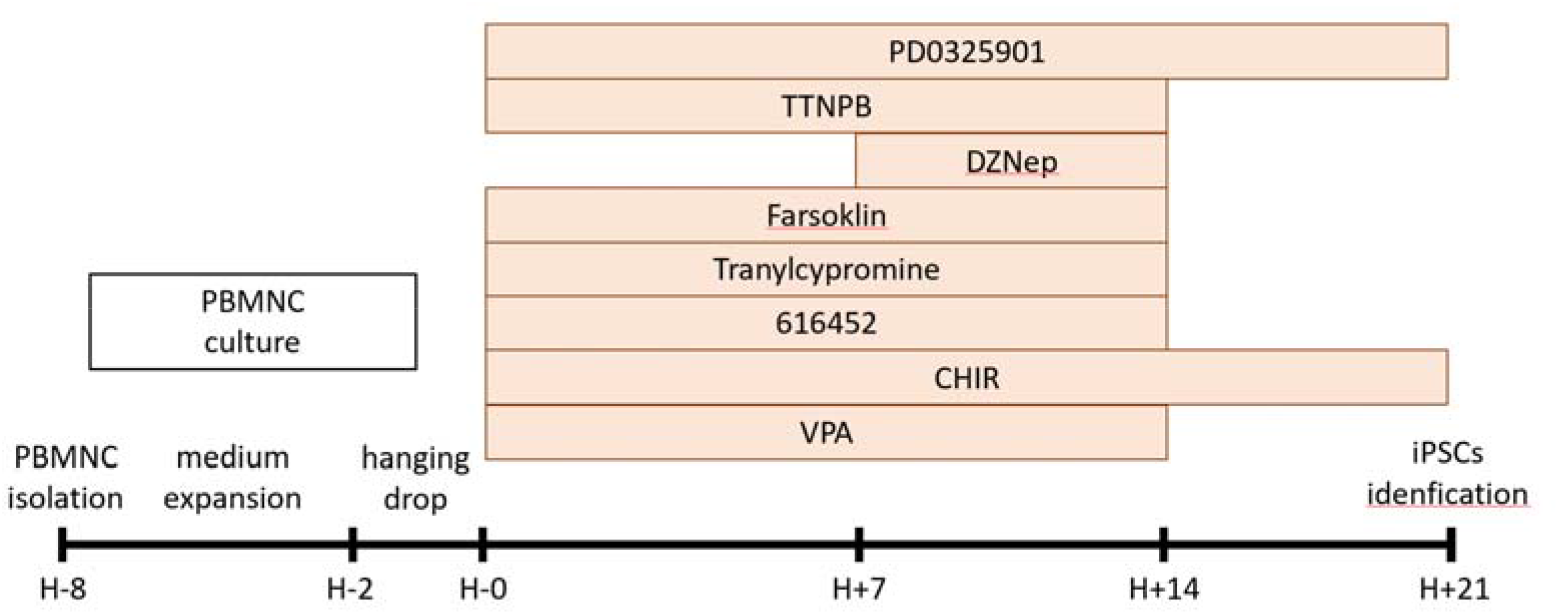
Study Protocol

### 2.1 PBMCs isolation

Blood was collected into heparin-coated tubes. Collected blood was diluted with 5 cc phosphate-buffered saline (PBS) and centrifuged through a Ficoll gradient for 30 minutes at 1600×g. PBMCs were collected and transferred to a new tube and add 10 cc PBS, centrifuged for 5 minutes at 2000-2500×g. Supernatant was aspirated and resuspend at 10 cc PBS.

### 2.2 Mononuclear cell reprogramming using Small Molecule Compound VC6TFZ

Three drops of mononuclear cell suspension were transferred to M-24 well plate pre-coated with vitronectin overnight and feeder cells. Feeder cells were made from mitosis-inactivated rabbit adipose mesenchymal cells using mitomycin-C 2 μg/mL for 30 minutes. We add ReproTesR medium which contained small molecule VC6TF. At day 7, DZnep was added. Cells were exposed to small molecules for 14 days. The medium was changed every 4 days. On day 14, VC6TFZ was changed to 2i medium (PD0325901 and CHIR).

Mononuclear cells were divided into 4 groups consisting of the control group (P1), and the experimental group (P2, P3, and P4) who received the small molecule compound VC6TFZ with different dosage variations; such as:

- Group 2 (P2) was the dosage VPA 0.5 mM, CHIR 5 μM, 616452 5 μM, tranylcypromine 2.5 μM, FSK 20 μM, DZnep 20 μM, and TTNPB 5 μM.
- Group 3 (P3) was the experimental groups which exposed to small-molecule dosage VPA 0.75 mM, CHIR 10 μM, 616452 7.5 μM, tranylcypromine 5 μM, FSK 40 μM, DZnep 50 nM, and TTNPB 5 μM.
- Group 4 (P4) was the experimental groups which exposed to small-molecule VPA 1 mM, CHIR 20 μM, 616452 10 μM, tranylcypromine 10 μM, FSK 60 μM, DZnep 100 μM, and TTNPB 5 μM.

### 2.3 PBMCs identification

Identification of iPSCs was done by morphology identification and expression of pluripotency markers (SSEA4 and TRA 1-60). Induced pluripotent stem cells colonies had large characteristics, tight and clear border, and cobble stone-like appearance. In iPSCs colonies, cells had a small size with a large ratio of cytoplasmic nuclei.

### 2.4 Immunocytochemical staining

Cells were fixed with methanol for 10 minutes, then wash with cold PBS 2 times. The cells were then stabilized with 0.25% Triton X-100 on PBS for 10 minutes at room temperature. The cells were washed with PBS 3 times for 5 minutes. Cells were incubated with 1% bovine serum albumin (BSA) in phosphate-buffered saline with Tween (PBST) for 30 minutes. Cells were then incubated in antibody solution (SSEA4 and TRA1-60) in 1% BSA in PBST in humidified chamber for 1 hour at room temperature or overnight at 4°C. The procedure continued with washing the cells for 5 minutes with PBS, repeated for 3 times. The cell was then incubated with the Alexa Fluor 488 goat anti-mouse IgG secondary antibody. Fluorescent signal is then viewed under a microscope.

### 2.5 Statistical analysis

Collected data were coded, tabulated, and statistically analyzed using the SPSS version 24. SSEA4 and TRA 1-60 expressions will be presented mean ± SD. Data normality test was done using Kolmogorov–Smirnov test. Differences in SSEA4 expression and TRA1-60 expression between the four groups will be analyzed by one-way ANOVA test if the data were normally distributed and Kruskal–Wallis if the data were not normally distributed. Significance cut off value is α = 0.05.

## 3. Results

### 3.1 Isolation and Culture of Peripheral Blood Mononuclear Cells

The isolation process aims to separate mononuclear cells from other cells in the peripheral blood such as erythrocytes and platelets. This isolation process is carried out by the centrifugation gradient density method where the mononuclear cells are in the buffy coat layer (Figure 3A). Mononuclear cell culture was carried out for 6 days by culturing it on basal medium (RPMI) enriched with L-Glutamine 1 mL / 100mL, ITS 1 mL / 100mL, FGF 5 ng / uL, ascorbid acid 5mg / 100mL, GMCSF 50uL / 100 mL, and dexamethasone 100uL / 100mL. The culture for 6 days aims to increase the efficiency of reprogramming.

**Figure 3.**
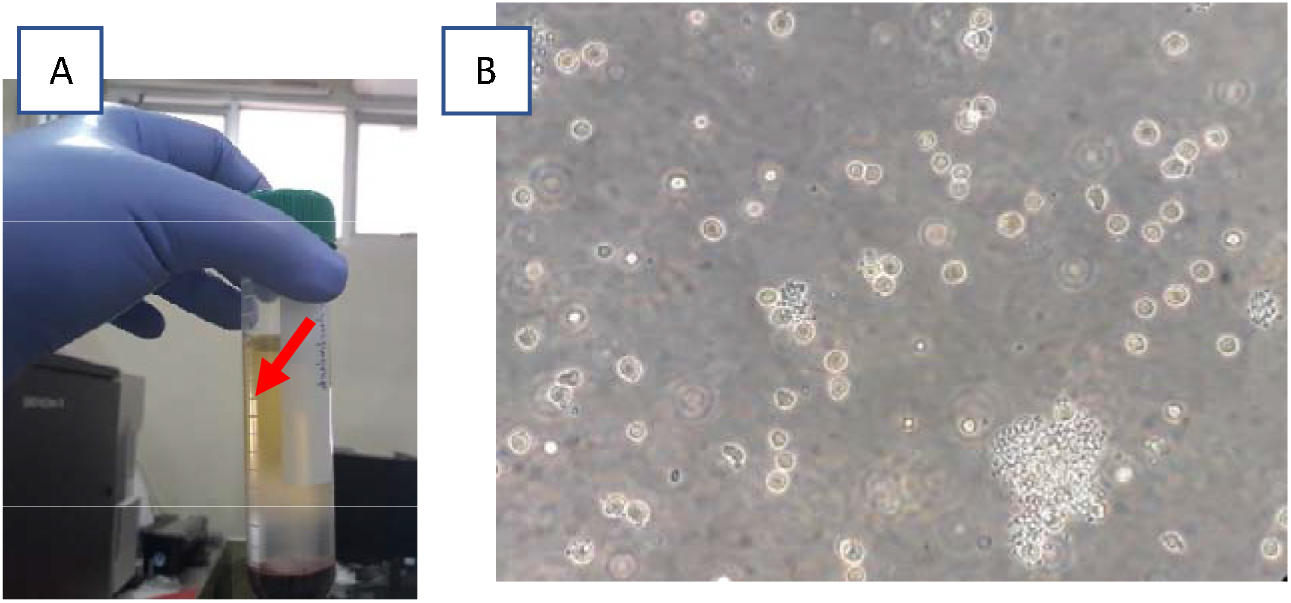
(A) Buffy coat layer between plasma and Ficoll, (B) Peripheral blood mononuclear cells (200x enlargement)

Mononuclear cells that had been cultured on medium enriched with cytokine cocktails for 6 days were then cultured using the hanging drop method for 48 hours (Figure 3B). Hanging drop is done by dripping mononuclear cells vertically 1 drop 25 μL of mononuclear cell suspension on a sterile petri-dish cover with a diameter of 5 cm and then turning it over (Figure 4A).

**Figure 4.**
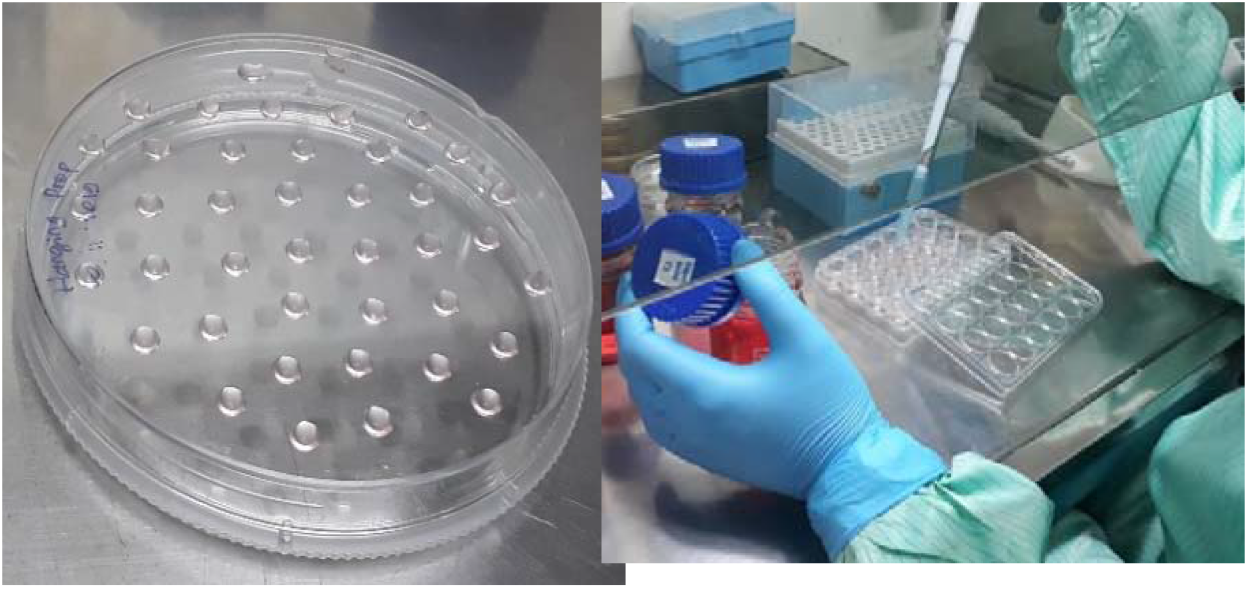
(A) Hanging drop method, (B) M6 well plate which was previously coated with vitronectin and feeder cell

### 3.2 Induction of Peripheral Blood Monouclear Cell Pluripotency

Five drops of mononuclear cell suspension that had been cultured by hanging drop method were plated on M-6 well plates that had been previously coated with vitronectin and feeder cells (Figure 4B). In this study, adipose mesenchymal stem cells (AMSC) were used in rabbits that had been pre-treated with 2 ug / mL of Mytomycin C (MMC).

The induction of munonuclear cell pluripotency was carried out for 21 days. During the first 7 days the cells received small molecule VPA, CHIR, 616452, Tranylcypromin, Farsocline, and TTNPB. Deazaneplanocin (DZnep) was added on day 7 to day 14. On day 14 the small molecule compound VC6TFZ was replaced with 2i medium consisting of PD0325901 and CHIR. ReproTesr Medium is changed every 4 days.

### 3.3 Identification of Induced Pluripotent Stem Cells

The identification of iPSCs in this study was carried out by identifying pluripotent stem cell colonies and the expression of markers SSEA 4 & TRA 1-60. The induced pluripotent stem cells tend to form colonies with large size, clear and clear edges. With the hanging drop method, it is possible to form colonies from the first day of plating. Colonies with large and round morphology, cobble stone-like appearances, and clear boundaries have been seen in 7th day of the reprogramming process (Figure 5).

**Figure 5.**
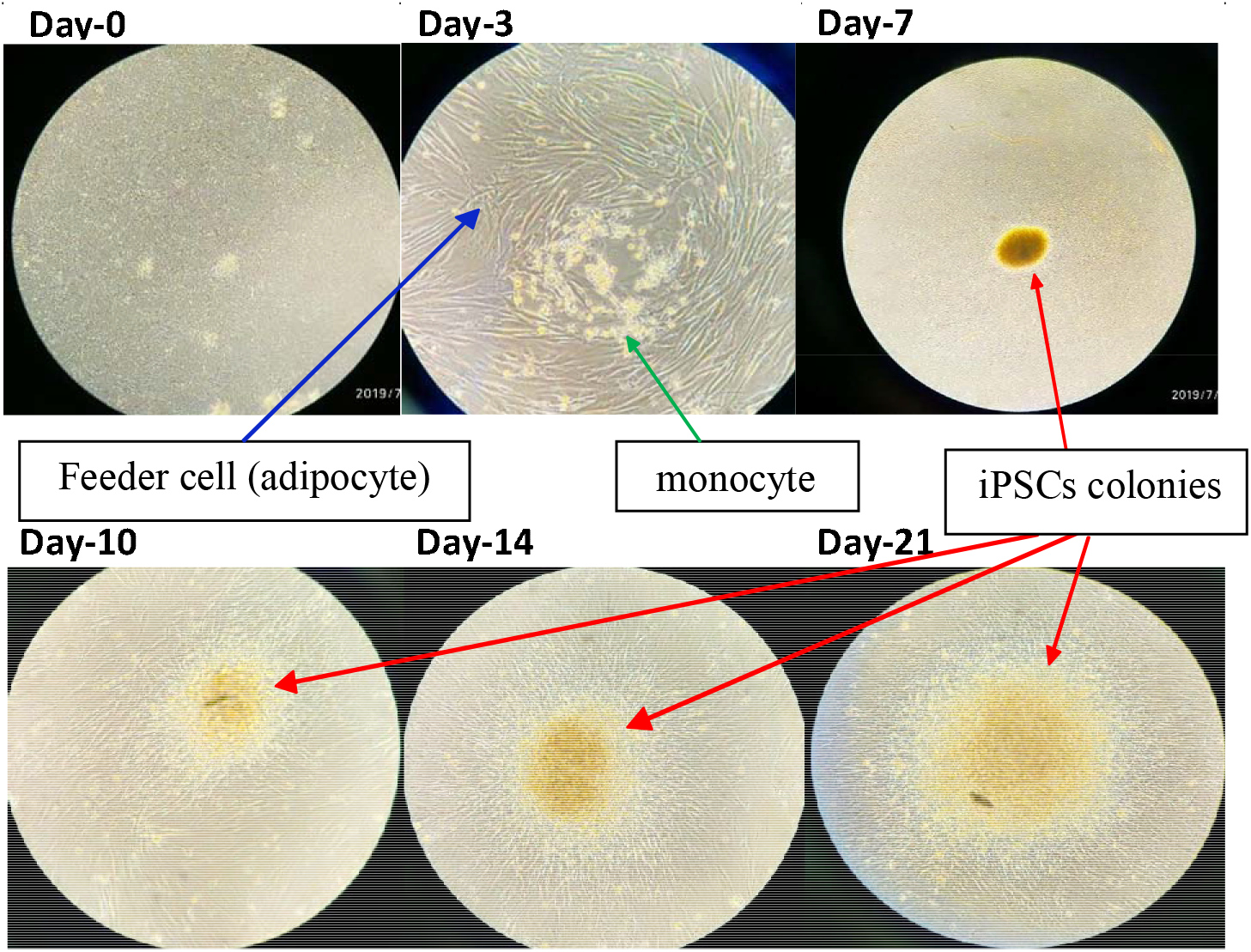
Changed of colonies morphology at day 0 until day 21

All colonies were expressed pluripotency markers. SSEA4 expression was found in all groups with the strongest expression in P3 group. Cells that had strong expressions of SSEA4 will glow bright green (Figure 6), the degree of luminance will be measured quantitatively using ImageJ software. From Kolmogorov-Smirnov, the quantitative data of SSEA 4 marker expression at each dose was normally distributed. Significant differences were found between all experimental groups (P2, P3, and P4 respectively) compared with the control group (p = 0.007, p = 0.001, p = 0.009), as shown in Table 1.

**Figure 6.**
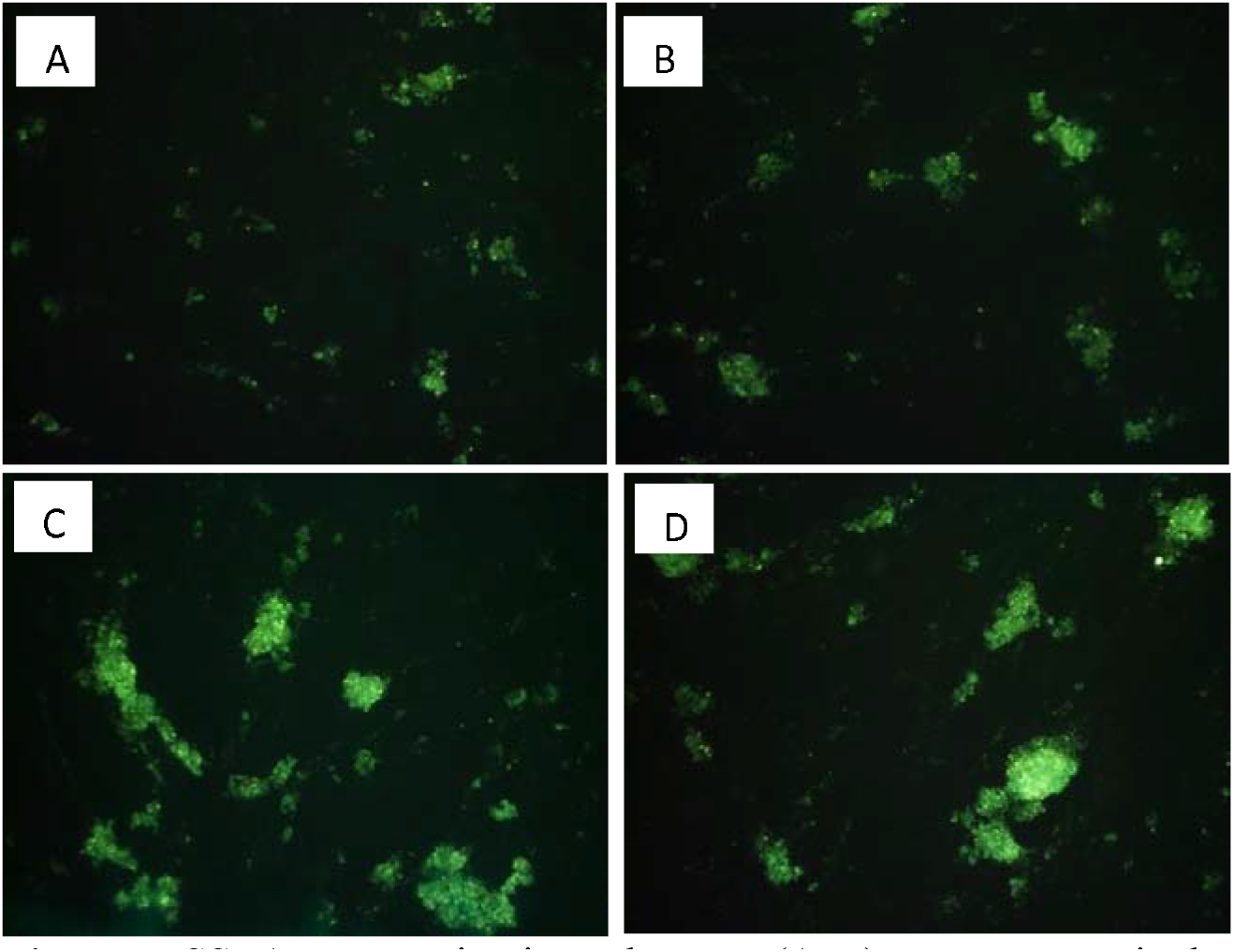
SSEA4 expression in each group (A-D) P1-P4 respectively

**Table 1.**
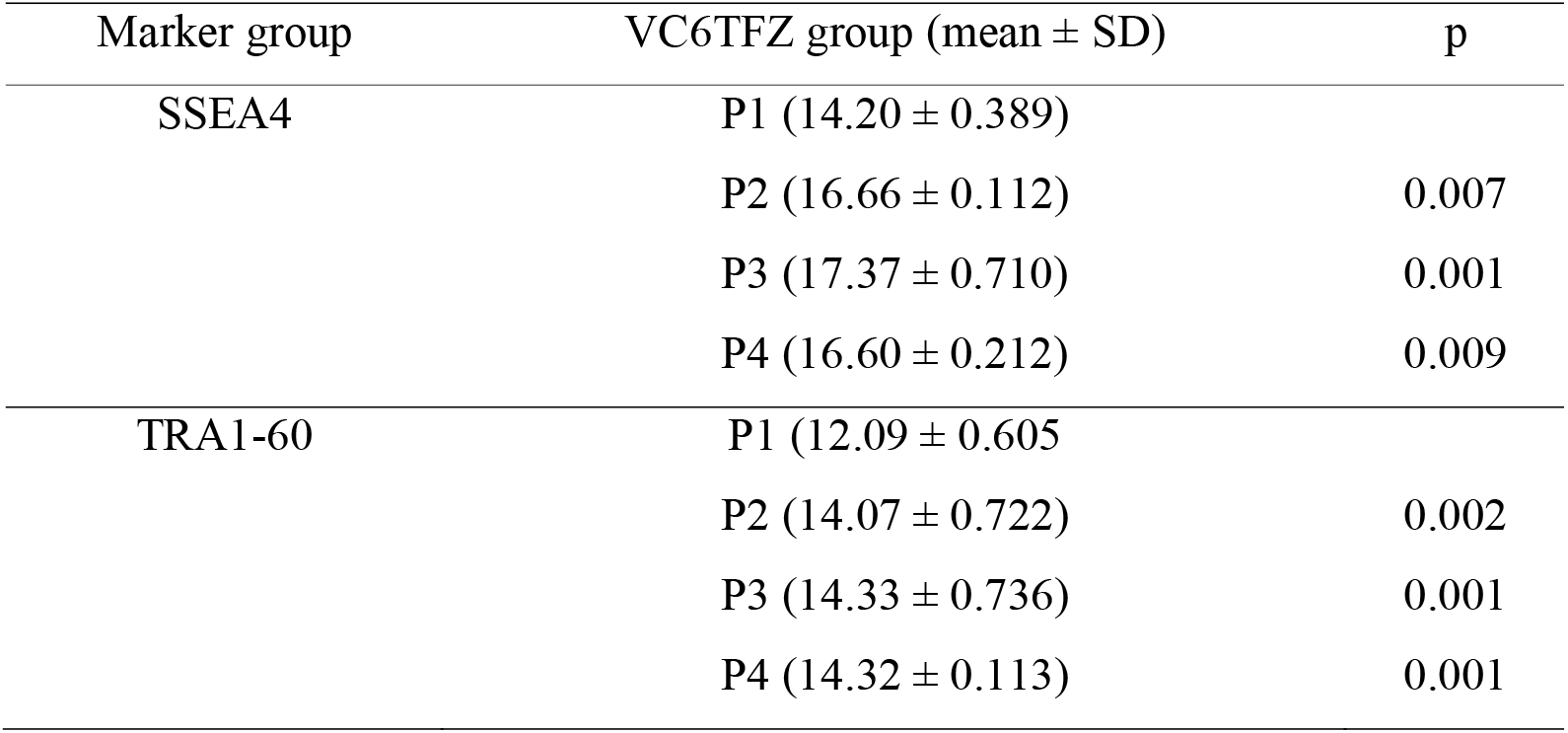
Comparison of expression between groups SSEA4 and TRA1-60

TRA1-60 expression was found in all treatment groups with the strongest expression in P3 and P4 group. Cells that had strong expressions of SSEA4 will glow bright green (Figure 7). The quantitative data of TRA1-60 marker expression at each dose was normally distributed. There were significant differences were found between all experimental groups (P2, P3, and P4 respectively) compared with the control group (p = 0.002, p = 0.001, p = 0.001), as shown in Table 1.

**Figure 7.**
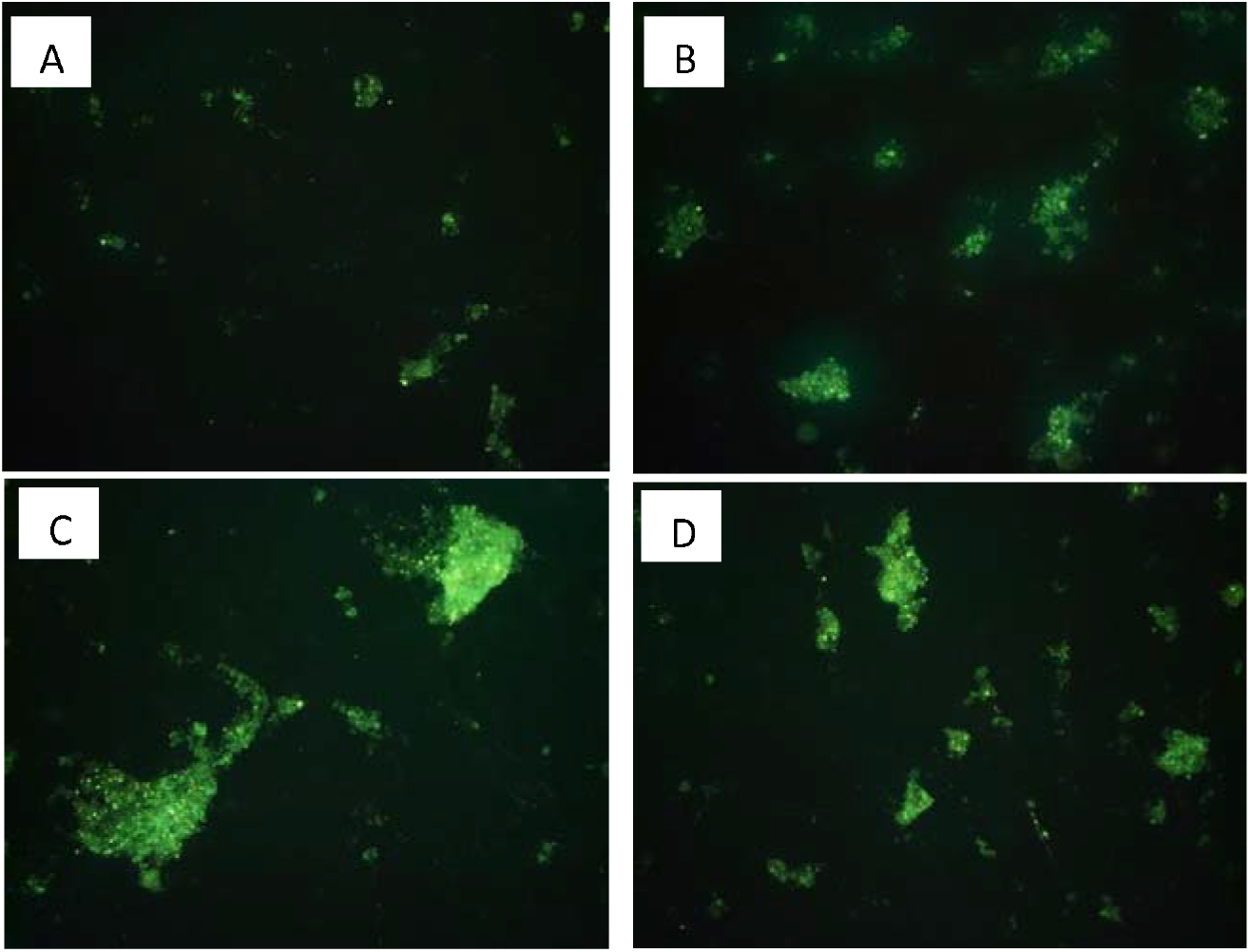
TRA1-60 expression in each group (A-D) P1-P4 respectively

## 4. Discussion

In this study, we found that small-molecule compound VC6TFZ could induce pluripotency in human PBMC. It was proven by the formation of colonies that resembled iPSCs morphology and the expression of pluripotency markers, SSEA4 and TRA1-60. In this study, we found a colony that began to appear on the 7th day of induction and increasingly enlarge on day 10. The colonies had round morphology, cobble stone like, and clear edges. This colonies morphology resembles iPSCs as described in the guidelines and techniques for the generation of iPSCs [7].

The colonies in this study express pluripotency markers, SSEA4 and TRA1-60. The expressions of each marker were analyzed quantitatively using ImageJ software, which obtained higher expressions in all treatment groups compared to the control group. Statistically, there were significant differences between the control group and the P3 and P4 groups in SSEA4 expression (p = 0,001; p = 0.009) and TRA1-60 (p = 0.001; p = 0.001). This indicates the most optimal expression of SSEA4 and TRA1-60 at small-molecule P3 and P4. This indicates that the effect of small molecule on the success of reprogramming was dose dependent [6,8]. The key for reprogramming successful with small molecules was the concentration and combination of small molecules. Small molecules could be cytotoxic at certain concentrations so that at higher doses, the reprogramming efficiency was sometimes even lower [8].

The protocol we used in this study for the induction was done in 21 days, faster than the Hou *et al.* In Hou’s study, GFP-positive colonies began to appear around day 10–12, while in this study, the colonies appeared earlier at day 6. This was likely due to the optimized culture method and the hanging drop culture method. Optimization of PBMC culture plays an important role in the reprogramming process. In this study, we cultured PBMC for 6 days in an expansion medium. During culture period, the longer duration, then greater number of dead cells, but the number of living cells remains constant, so the optimal duration of culture time was needed. This 6-day duration was based on research conducted by Gu *et al.* where this study showed that the 6-day culture time shows the number of colonies that most expressed TRA 1-60 and AP compared to days 4, 8 or 10 [9].

Hanging drop culture method allows the formation of colonies through cell aggregation induced by gravity [10]. The accumulation of cells in the drop allows the formation of spheroidal colonies. Spheroid colonies that were formed can produce extracellular matrices and environments that resemble living tissue. Inside the drop cells adhere to each other by holding on the resulting matrix. Intercell interactions and interactions between cells and extracellular matrix were better with this method [10]. This method also allows more efficient diffusion of growth factors and metabolic waste disposal [11].

One of the challenges in reprogramming PBMC was that these cells had non-adherent nature [12]. To overcome this problem, we coated the well walls using vitronectin and feeder cells that enable the PBMC attachment on the well. This attachment was important to avoid cell lost during medium replacement. This feeder cell produced adhesion molecules and extracellular matrix that increases attachment of iPSCs which supports growth and survival [12,13].

The quantitative data analysis in this study was based on the results of the immunocytochemistry (ICC) test, which is a common laboratory examination technique used to anatomically visualize the localization of proteins or specific antigens in cells using specific primary antibodies that bind to these proteins. In this study, the method used for checking ICC is the indirect method. This method has a disadvantage, namely that secondary antibodies can cross-react with proteins other than the target, so that it can provide a luminescence image that is not specific to the target cells. However, the advantage of this method is that it costs less, because it can be done by using the same conjugated secondary antibody to detect different primary antibodies [14].

## 5. Conclusions

Small molecule compound VC6TFZ could induced pluripotency of human PBMC to generate iPSCs. Pluripotency marker gene expression, SSEA 4 and TRA 1-60, in the experimental group was statistically significantly higher than in the control group. Further research was needed to analyze molecular profile, differentiation, and self-renewal ability of these cells. For clinical application, the safety profile related to the risk of teratogenesis and genetic instability also needed further investigation.

## Acknowledgements

Not applicable

